# Five years of GenoTyphi: updates to the global *Salmonella* Typhi genotyping framework

**DOI:** 10.1101/2021.04.28.441766

**Authors:** Zoe A. Dyson, Kathryn E. Holt

## Abstract

In 2016 a whole genome sequence (WGS) based genotyping framework (GenoTyphi) was developed providing a phylogenetically informative nomenclature for lineages of *Salmonella* Typhi, the aetiological agent of typhoid fever. Subsequent surveillance studies have revealed additional epidemiologically important subpopulations, necessitating the definition of new genotypes and extension of associated software to facilitate the detection of antimicrobial resistance (AMR) mutations. Analysis of 4,632 WGS provide an updated overview of the global *S.* Typhi population structure and genotyping framework, revealing the widespread nature of H58 (4.3.1) genotypes and the diverse range of genotypes carrying AMR mutations.

## Background

Typhoid fever is a faeco-orally transmitted systemic infection caused by the bacterium *Salmonella* Typhi (*S.* Typhi). Each year >10 million cases occur worldwide of which >100,000 are associated with mortalities^1^ making it a public health threat in many low- to middle-income countries with limited hygiene and sanitation infrastructure.

*S*. Typhi is a genetically monomorphic pathogen with a slow mutation rate that infrequently recombines^2^. Whole genome sequencing (WGS) and core-genome phylogenetics have become the standard for typhoid molecular epidemiology in both research and public health settings, providing insights into population structure, transmission dynamics, AMR emergence and dissemination, as well as outbreak investigation and monitoring of implemented intervention strategies. In 2016 a WGS based genotyping framework for *S.* Typhi was developed using a collection of ~2,000 genomes from >60 countries^3^, with the goal of stratifying the pathogen population and providing a phylogenetically informative nomenclature with which to refer to different lineages. The resulting scheme (known as GenoTyphi) utilised 68 marker single nucleotide variants (SNVs) to define, based on an inferred genome-wide phylogeny, four primary clades, 16 clades, and 49 subclades organised in a pseudo-hierarchical nomenclature whereby primary clade 1 is subdivided into clades 1.1 and 1.2; clade 1.1 is further subdivided into subclades 1.1.1, 1.1.2, and 1.1.3. Haplotype 58 (H58), which has previously been associated with antimicrobial resistance (AMR) and global dissemination via intercontinental transmission events^2^, was designated genotype 4.3.1 under the new scheme. A software tool for calling GenoTyphi genotypes from WGS data was implemented in Python (available at: https://github.com/katholt/genotyphi), facilitating integration of the scheme into bioinformatics pipelines. GenoTyphi is also available to non-expert users via the online data analysis platform Typhi Pathogenwatch (https://pathogen.watch/)^4^.

Following publication of the initial framework, regional surveillance studies identified additional epidemiologically important subpopulations of *S*. Typhi, necessitating definition of new genotypes^5–9^. Further, point mutations responsible for reduced susceptibility to fluoroquinolones and azithromycin have also emerged^10,11^, necessitating extension of the GenoTyphi software tool for their detection. Here we provide an overview of updates to both the GenoTyphi scheme and pipeline (summarised in **Tables S1-S2**), as well as the view it provides of the global pathogen population.

## Materials and methods

### Phylogenetic and SNV analysis of S. Typhi isolates

Reads from 4,632 *S.* Typhi genomes (**Table S3**) were mapped to the reference sequence of *S.* Typhi CT18 (accession number: AL513382) with RedDog (vbeta.11; available at: https://github.com/katholt/RedDog). Sequences were assigned to genotypes, and quinolone resistance determining region (QRDR) and *acrB* mutations associated with AMR identified detected, using GenoTyphi (v1.9.1; available at: https://github.com/katholt/genotyphi) which is permanently archived by Zenodo at doi: 10.5281/zenodo.4707614. Recombinant regions were removed from the whole genome Single Nucleotide Variant (SNV) alignment using Gubbins (v2.4.1; available at: https://github.com/sanger-pathogens/gubbins) and a maximum-likelihood phylogeny inferred with RAxML (v8.2.9; available at https://github.com/stamatak/standard-RAxML). An interactive annotated phylogeny is available at https://microreact.org/project/vBoskUuenEVmfVzrcAMx8R. Further details are provided in **supplementary methods**.

## Results

### Global overview of S. Typhi genotypes

Analysis of 4,632 published genomes demonstrate that H58 has now disseminated across most continents (**Fig. 1a**), with the distribution of genotypes differing per country (**Fig. 1b**). The 82 genotypes defined at present (**Fig. 1c; Table S1**) include those from the original publication, subdivision of 4.3.1 (H58) into three major lineages (4.3.1.1, 4.3.1.2, 4.3.1.3), genotypes designating newly identified subclades (e.g. 2.5.2, 3.3.2), and designations for AMR populations of epidemiological importance (e.g. 4.3.1.1.P1).

**Figure 1.**
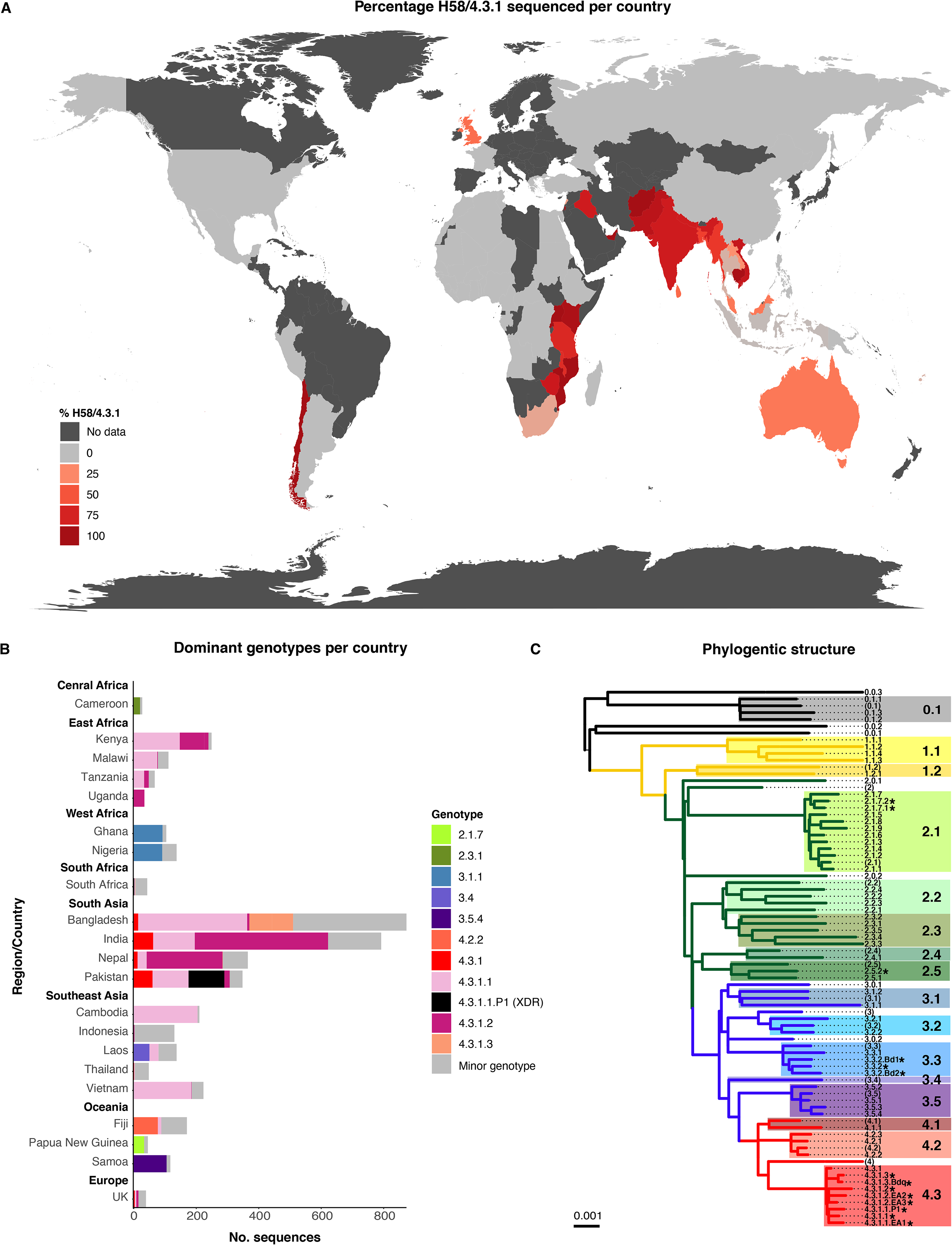
Global genotype distribution and population structure. **(A) Global dissemination of genotype 4.3.1 (H58).** Countries are coloured by total percentage of H58 genotypes amongst isolates in the genome collection, as per inset legend. **(B) Dominant genotypes per location.** Dominant genotypes (each accounting for >30% of sequenced isolates per country) and H58 genotypes are coloured as per the inset legend, with minor non-H58 genotypes in grey. Genotypes are shown for countries with at least 20 genome sequences. **(C) Phylogenetic tree backbone showing the relationships between 16 clades and 63 subclades/sublineages. Tree tips represent unique genotypes as labelled, background shading highlights clades (labelled in larger font).** * indicates genotypes added to the scheme following its initial publication, brackets indicate undifferentiated clades and primary clades.

### Updated H58 (4.3.1) genotypes

Genotype 4.3.1 is currently subdivided into three lineages (see **Fig. 1c**; **Table S1**). H58 lineages I (genotype 4.3.1.1) and II (genotype 4.3.1.2) were originally defined in a study of paediatric patients attending Patan Hospital in Kathmandu, Nepal^12^. Later studies^8^ revealed the co-circulation of both lineages in this setting between 2008-2016, with a shift in dominance to 4.3.1.2 after 2010 (40% 4.3.1.2 pre-2010 and 74% post-2011, p=1.0×10^7^), warranting more discriminant typing to capture such changes in population structure. H58 lineage III (genotype 4.3.1.3), originally defined in an examination of 536 AMR sequences from Dhaka, Bangladesh^6^, is a monophyletic cluster of genotype 4.3.1 mostly from Bangladesh (99%). It was recently detected at a frequency of 9% in urban Dhaka between 2004-2016^5^. A monophyletic sublineage of genotype 4.3.1.3 was resistant to fluoroquinolones (median minimal inhibitory concentration (MIC) of 4 μg/ml) and is here formally designated 4.3.1.3.Bdq based on previous studies^6^.

A recent study of asymptomatic carriers and acute typhoid fever patients in Kenya detected the co-circulation of genotypes 4.3.1.1 and 4.3.1.2^9^. Contextualisation with the global phylogeny attributed the presence of these lineages to two previously reported transmission waves originating in South Asia^2,13,^ and a third more recent introduction of 4.3.1.2 from South Asia that has apparently also reached Uganda^9^. These three H58 sublineages were each comprised exclusively East African sequences and had different AMR profiles, and thus were designated as new genotypes in order to help monitor their spread: H58 lineage I sublineage East Africa I (4.3.1.1.EA1), H58 lineage II sublineage East Africa II (4.3.1.2.EA2) and H58 lineage II sublineage East Africa III (4.3.1.2.EA3; **Fig. 1c**; **Table S1**)^9^.

In 2016, outbreaks of the first widespread extensively drug resistant (XDR) clone occurred in Pakistan. This monophyletic outbreak cluster of genotype 4.3.1.1, resistant to chloramphenicol, ampicillin, and co-trimoxazole, fluroquinolones and third generation cephalosporins^7,14^, was designated genotype 4.3.1.1.P1 to aid its identification.

### Updated non-H58 genotypes

Studies of *S.* Typhi in Bangladesh^5^ revealed 119 genomes (14.5% of sequences analysed) formed a monophyletic group of sequences typed only to clade level (genotype 3.3) that were related to sequences from Nepal (separated by ~70 SNVs, also typed as 3.3). These were collectively designated genotype 3.3.2. Within the Bangladesh 3.3.2, two sublineages carrying QRDR mutations were further defined to facilitate their detection in future surveillance studies; 3.3.2.Bd1 (which typically carry *gyrA*-S83F), and 3.3.2.Bd2 (which typically carry *gyrA*-S87N) (see **Fig. 1c**; **Table S1**).

Ongoing analysis of genomes from Madagascar and Papua New Guinea (to be described in detail elsewhere) have also identified localised variants. The Madagascar group belongs to clade 2.5, is distantly related to other 2.5 sequences from India (separated by ~122 SNVs) and has been designated 2.5.2. The PNG genotype 2.1.7 population is subdivided into two distinct sublineages designated genotypes 2.1.7.1 and 2.1.7.2, with 2.1.7.1 observed more frequently.

### Updated detection of resistance-associated mutations

Aforementioned studies of paediatric typhoid in Kathmandu, Nepal revealed high levels (75.3%) of sequences carrying non-synonymous point mutations in the QRDR of genes *gyrA, gyrB,* and *parC* responsible for reduced susceptibility to fluoroquinolones from 2008-2016^8^. Among these were sequences of genotype 4.3.1.2 carrying three such mutations (e.g. *gyrA*-S83F, *gyrA*-D87N, *parC*-S80I – 7.6%; *gyrA*-S83F, *gyrA*-D87N, *parC*-E84K – 0.5%) the former of which was previously found to cause treatment failure among adult populations in the same setting^11^. More recent studies^10^ demonstrated that mutations at codon 717 of gene *acrB*, a component of the AcrAB-TolC drug efflux pump, mediate Azithromycin resistance (MIC ≥32 μg/ml) in *S.* Typhi and had been observed at low frequency in Dhaka, Bangladesh (~1.3% of all *S.* Typhi isolated from 2009-2016). Subsequently, the GenoTyphi pipeline has been extended to detect mutations in both the QRDR and codon 717 of gene *acrB* (see **Table S2**).

### Global overview of AMR associated mutations

Analysis of published genomes demonstrates that sequences carrying QRDR mutations can now be found across most continents (**Fig. 2a**), with the diversity of genotypes carrying QRDR mutations varying by geographic region (**Fig. 2b**). The geographic distribution of sequences carrying *acrB*-R717Q/L mutations associated with azithromycin resistance are shown in **Fig. 2a-b**. These mutations have emerged independently in multiple *S.* Typhi genotypes in several different countries, mostly in South Asia at present, and are accompanied by QRDR mutations making them co-resistant to fluoroquinolones (see **Fig. 2b**). Isolates from Dhaka^10^ have also been reported to be multi-drug resistant carrying genes conferring additional resistance to chloramphenicol, ampicillin and co-trimoxazole. Recent studies^15^ has revealed that these mutations now appear to be emerging in more non-H58 genotypes in Dhaka from 2016 onwards including genotypes 2.3.3, 3.2.2, and 3.3.2.

**Figure 2.**
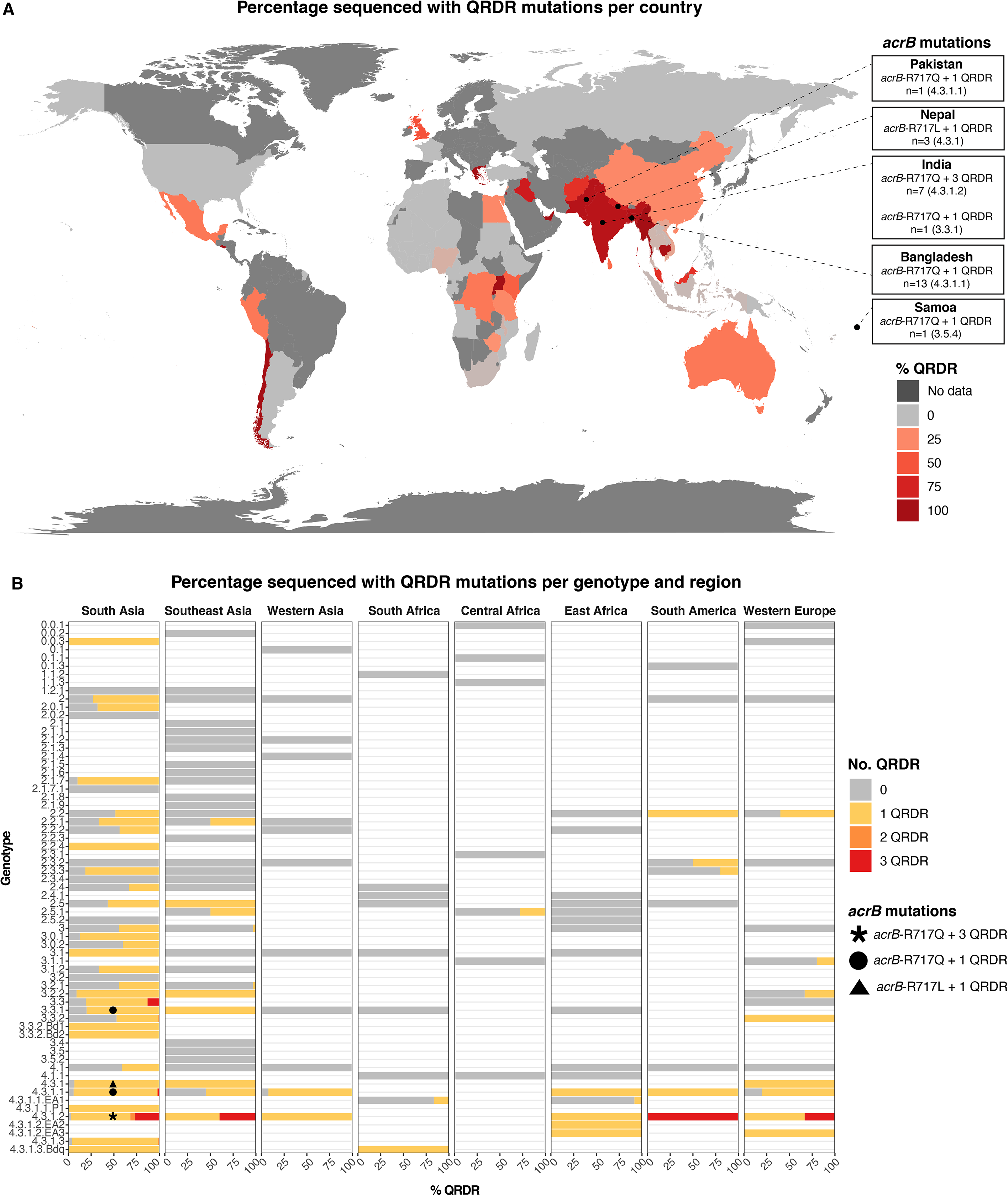
Global overview of AMR mutations. **(A) Global distribution of QRDR mutations.** Countries are coloured by the total percentage of sequences carrying QRDR mutations in the genome collection, as per inset legend. Locations where sequences also carrying acrB-R717Q/L mutations have been isolated are indicated as labelled. **(B) Distribution of QRDR mutations by genotype and region.** Genotype/region combinations are included where >25 isolates have been sequenced from the region and >5% of those carry QRDR mutations. Genotypes also carrying acrB-R717Q/L mutations are labelled as per the inset legend.

## Discussion

In the five years since the publication of the GenoTyphi framework several regional genomic surveillance studies have been carried out, providing further insight into transmission events on a regional and global scale (including the continued global spread of 4.3.1 genotypes, and the emergence, spread, and ongoing evolution of mutations responsible for AMR in a diverse range of H58 and non-H58 genotypes) and the identification of new genotypes (**Tables S1-S2**).

The GenoTyphi framework will continue to be developed as new data becomes available and as new variants emerge, providing up to date phylogenetically informative nomenclature for identifying and discussing trends in population structure and evolution of AMR in *S.* Typhi. This nomenclature remains critical in genetic epidemiology studies required for the successful implementation and monitoring of control strategies. Requests for the inclusion of new genotypes can be made via the GitHub repository (https://github.com/katholt/genotyphi), and will be overseen by the Global Typhoid Genomics Consortium steering committee (https://www.typhoidgenomics.org/).

## Supplementary data

- ***Supplementary methods***
- ***Table S1 - Summary of S. Typhi genotypes (Excel spreadsheet).** ‘Reference allele’ indicates the allele in the CT18 reference sequence. ‘Alternative allele’ indicates an allele called against the CT18 reference sequence for the genotype called. ‘Derived allele’ indicates the subtree-defining allele, which resulted from mutation of the original (ancestral) allele at this position to generate a new (derived) allele that we use as the marker for the subtree that corresponds to this genotype. ‘Ancestral allele’ indicates the allele present in the ancestor of S. Typhi, which is conserved by all members of the population outside of the subtree that corresponds to this genotype.*
- ***Table S2 - Summary of S. Typhi AMR mutations detected by GenoTyphi (Excel spreadsheet). ‘**Reference allele’ indicates the allele in the CT18 reference sequence. ‘Alternative allele’ indicates an allele called against the CT18 reference sequence for the genotype called.*
- ***Table S3 – Details of publicly available S. Typhi genome sequences analysed in this study (Excel spreadsheet)***

## Presentation statement

This work was carried out for an invited presentation ‘A global genomic perspective on Typhoid and AMR’ by Zoe Anne Dyson at the 15^th^ Asian Conference On Diarrhoeal Disease and Nutrition (ASCODD) on the 30^th^ of January 2020 in Dhaka, Bangladesh.

## Affiliation changes

The authors have not changed affiliations since this study was completed.

