## Supplementary methods for "Five years of GenoTyphi: updates to the global *Salmonella* Typhi genotyping framework"

***Phylogenetic and SNV analysis of S. Typhi isolates***

A total of 4,632 *S.* Typhi genomes published in various studies were included (accessions in **Table S3**)^1-18^. For Single Nucleotide Variant (SNV) analysis, paired-end reads were mapped to the reference sequence of *S.* Typhi CT18 (accession number: AL513382)^19^ using the RedDog mapping pipeline (vbeta.11), available at: <https://github.com/katholt/RedDog>. RedDog uses Bowtie (v2.2.3)^20^ to map reads to the reference sequence; SAMtools (v0.1.19)^21^ to identify SNVs with phred quality scores above 30; to filter out SNVs supported by <5 reads, or with 2.5 times the genome-wide average read depth (representing putative repeated sequences), or with ambiguous (heterozygous) consensus base calls. For every SNV position that passes the aforementioned criteria in any one isolate, consensus base calls (i.e. alleles) for that position were extracted from all genomes analysed, and used to construct an alignment of alleles across all SNV sites. Resultant read alignments (BAM format) were then used to assign sequences to previously defined genotypes (**Table S1**) according to the *S.* Typhi extended genotyping framework with the GenoTyphi (v1.9.1) pipeline (available at: <https://github.com/katholt/genotyphi>)^18^ which is permanently archived by Zenodo^22^ at doi: 10.5281/zenodo.4707614. AMR mutations in the quinolone resistance determining region (QRDR) and at codon 717 of gene *acrB* (detailed in **Table S2**) were detected in the same manner with GenoTyphi.

For phylogenetic analyses, chromosomal SNVs with confident homozygous calls (phred score >20) in more than 95% of the genomes mapped (representing a “soft” core genome) were concatenated to form an alignment of alleles at 36,859 variant sites, and alleles

from *S.* Paratyphi A str. AKU_12601^23^ (accession number: FM200053) were also included in the phylogenetic analysis as an outgroup for tree rooting. SNVs called in phage regions or repetitive sequences (354 kbp; ~7.4% of bases in the CT18 reference sequence, defined previously^5,18,24^) were filtered from the alignment, which was then used with the CT18 reference genome to produce a whole genome pseudoalignment that was subjected to recombination filtering with Gubbins (v2.4.1)^25^ to remove any further recombinant regions. This resulted in a final set of 35,599 SNVs identified in a final alignment length of 4,275,037 sites for the 4,633 sequences.

From the global SNV alignment, a maximum likelihood (ML) phylogenetic tree was inferred using RAxML (v8.2.9)^26^, with a generalised time-reversal model, a Gamma distribution to model site-specific rate variation (the GTR+ Γ substitution model; GTRGAMMA in RAxML), and 100 bootstrap pseudo-replicates to assess branch support.

Phylogenetic trees were visualised using Microreact^27^ and the R package ggtree (v2.2.4)^28^. An interactive phylogeny annotated with AMR mutations and genotypes is available at: <https://microreact.org/project/vBoskUuenEVmfVzrcAMx8R>. For the purpose of plotting the global genotype structure in **Fig. 1c**, clusters of *S.* Typhi in the full tree that were members of the same genotype were collapsed into a single representative each using the *drop.tip()* function in the R package ape^29^.
